# Increased expression of *Cd74* and MHC II genes by aortic macrophages links atherosclerosis with aging in mice

**DOI:** 10.64898/2026.07.17.739099

**Authors:** Adyasha Mishra, Inderjit S. Dhami, Shalini Kumari, Supriya Borah, Keerthana Goud, Shrinidhi Sahadevan, Balan Ramesh, Thierry Huby, Helen S. Goodridge, Pijus K. Barman

## Abstract

Atherosclerosis is a chronic inflammatory condition of the arteries leading to myocardial infarction, ischemic stroke and peripheral arterial diseases. Atherosclerotic plaques, which obstruct blood flow contain functionally diverse macrophage populations including pro-atherogenic and atheroprotective subsets. Aging is a major risk factor of atherosclerosis, but the mechanism underlying aging associated risk of atherosclerosis is unclear. Here, using integrated single-cell RNA sequencing data analysis, we demonstrate that specific monocyte and macrophage subsets are enriched in atherosclerotic plaques and the aging aorta of mice. We also show that *Cd74* and MHCII genes such as *H2-Aa, H2-Ab1, H2-Eb1*, and *H2-DMb1* are consistently upregulated in the monocytes and macrophages from atherosclerotic plaques, the aging aorta, and the aging bone marrow of mice. Our experimental data also show increased expression of CD74 surface protein by aortic macrophages and bone marrow monocytes from aged mice. In addition, we show increased CD74 expression by RAW264.7 mouse macrophages and THP1 human monocytes following ox-LDL stimulation in vitro. Finally, we demonstrate that monocytes from aged mice adhere more to aortic endothelial cells in co-cultures. Thus, increased monocyte adhesion to endothelial cells may explain the enhanced proportion of specific monocyte and macrophage subsets in the aging aorta, inducing a pre-atherosclerotic condition.

## Introduction

Aging is a major risk factor for atherosclerosis, which is characterized by monocyte arrest on vascular endothelial cells, their trans-endothelial migration and transformation into foamy macrophages by oxidized LDL (ox-LDL) uptake, and plaque formation ^1^. Various studies have revealed the presence of functionally diverse monocyte and macrophage subsets in atherosclerotic plaques ^2–8^. For instance, while some plaque macrophages are pro-atherogenic, others are believed to be atheroprotective ^3,5,9–13^. However, the origin of plaque-associated macrophages as well as the link between monocyte aging and plaque macrophage accumulation is unclear.

Monocytes are innate immune cells, which are produced by hematopoiesis in the bone marrow and infiltrate into peripheral tissues where they can differentiate into macrophages ^14–16^. Monocytes and macrophages play a variety of roles in anti-microbial defense, tissue repair, and antigen presentation ^17,18^. There are broadly three major subtypes of monocytes in humans and mice: i) “classical” monocytes (CD14^+^ CD16^−^ in humans and Ly6C^hi^ in mice), ii) “non-classical” monocytes (CD14^lo^ CD16^+^ in humans and Ly6C^lo^ in mice), and iii) “intermediate” monocytes (CD14^+^ CD16^+^ in humans and Ly6C^int^ in mice) ^16,19–21^. Furthermore, using multiparametric single-cell studies, we and others have revealed distinct classical monocyte subsets produced by granulocyte-monocyte progenitors (GMPs) and monocyte-dendritic cell progenitors (MDPs) in mouse bone marrow^17,22–26^.

Aging leads to a chronic low-level inflammation (inflammaging), which is associated with the increased production and altered function of monocytes ^27–32^. For instance, monocytes from aged humans show increased production of pro-inflammatory cytokine TNF-α ^33^, stronger adhesion to type I collagen ^34^, decreased mitochondrial respiration ^35^, and defective lipid metabolism ^36^. Monocytes and macrophages from aged mice display impaired pathogen clearance and defective phagocytosis of senescent neutrophils ^28,31,37^. In addition, we have previously demonstrated increased production of classical monocytes expressing CD74 and MHCII in aging mice ^24^. Importantly, our data revealed that MDPs and classical monocytes from the bone marrow of aged mice yield more monocyte-derived dendritic cells (MoDC; CD11c^+^ MHCII^+^ cells) in GM-CSF cultures. However, the impact of altered monocyte functions on atherosclerotic risk in aging is unclear. The current study aimed to fill this gap in the literature using integrated single-cell RNA sequencing (scRNA-seq) data analysis of monocytes and macrophages from atherosclerotic and aging mice, and experimental validation in aging mice and in vitro atheroma models.

Integrated scRNA-seq data analysis revealed that specific plaque-associated macrophages are also enriched in aging mouse aorta. Differentially expressed gene (DEG) analysis showed that *Cd74* and MHCII genes such as *H2-Aa, H2-Ab1, H2-Eb1*, and *H2-DMb1* are consistently upregulated in the monocytes and macrophages from atherosclerotic plaques, aged aorta, and aged bone marrow. Consistent with this, flow cytometry analysis showed that aortic macrophages and bone marrow monocytes from aged mice (26-month-old) have increased expression of CD74. In addition, RAW264.7 mouse macrophages and THP1 human monocytes showed increased CD74 expression after ox-LDL stimulation in vitro. Lastly, we observed that a higher number of bone marrow monocytes from aged mice adhered to aortic endothelial cells (AECs) in co-cultures. Together, these data may imply that aging increases monocyte adhesion to AECs, leading to recruitment of CD74^+^ monocytes to the aorta, which is a key feature of atherosclerosis.

## Results

### Specific monocyte and macrophage subsets are enriched in atherosclerotic plaques and aging aorta

We performed integrated scRNA-seq analysis of mouse aortic leukocytes (CD45^+^ cells) ^5^, aortic non-smooth muscle cells ^4^, aortic myeloid cells (CD11b^+^ cells) ^3^, total aortic cells ^38^, and bone marrow classical monocytes (Ly6C^hi^) ^24^ from atherosclerotic plaques and different aging conditions to examine atherosclerosis- and aging-associated alterations in cellular subsets (Table 1). We analyzed a final number of 45680 cells, and unsupervised uniform manifold approximation and projection (UMAP) clustering showed 33 clusters which included aortic tissue structural cells (11 clusters), lymphoid cells (8 clusters), monocytes and macrophages (9 clusters), dendritic cells (DCs) (3 clusters), neutrophils (1 cluster), and mixed or non-aortic cells (1 cluster) (Figure S1A, S1B and S2, and Table S1A and S2A). Study-wise segregation of UMAP plots revealed the contribution of each study to the cell types as anticipated (Figure S1C). Together, integrated scRNA-seq analysis showed consistent clustering and cell type identification as reported previously.

**Table 1.**
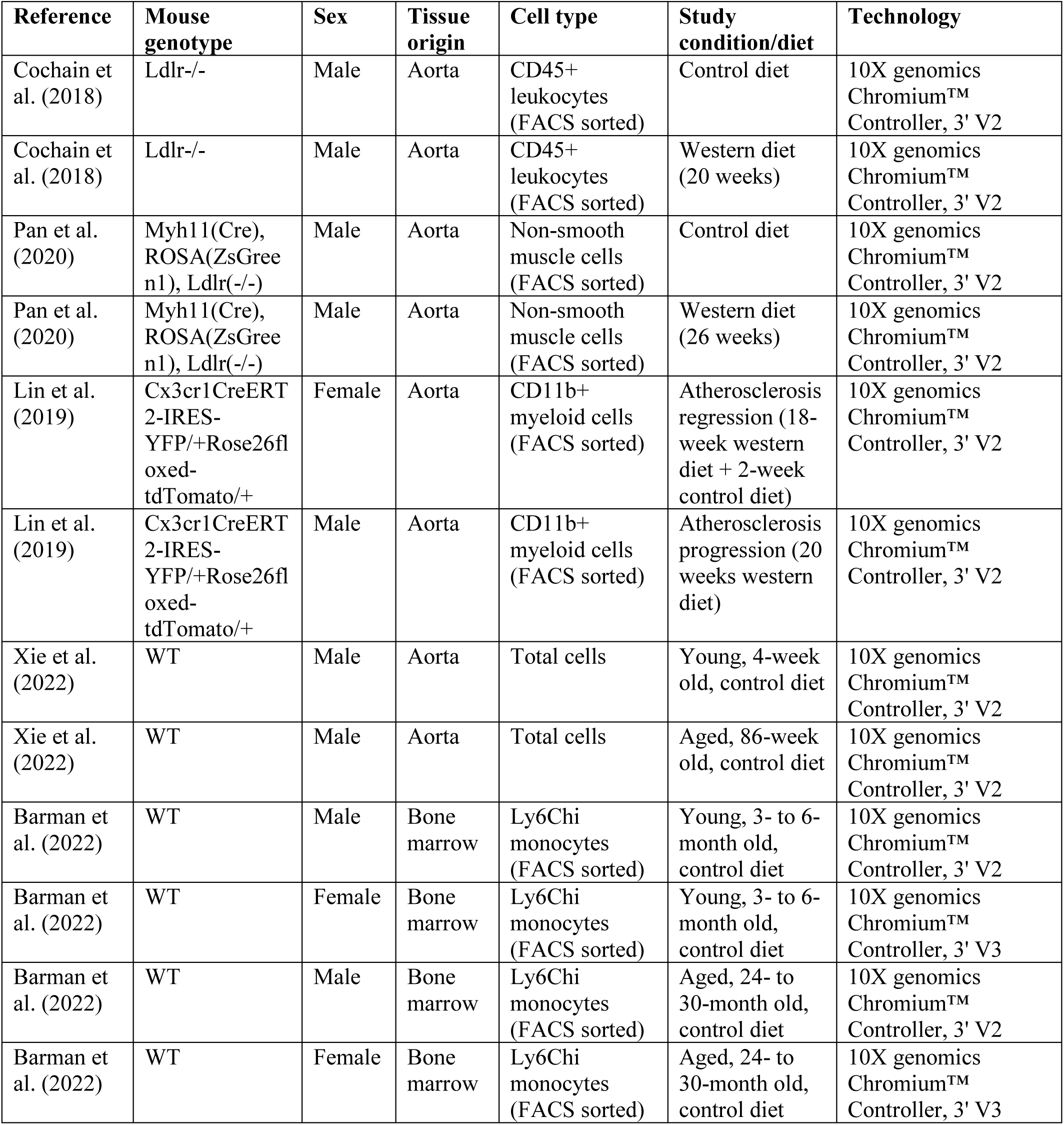
Mouse scRNA-seq datasets used in the study.

Next, we focused our analysis on monocytes and macrophages to examine association between bone marrow monocytes and aortic macrophages from atherosclerotic and aged mice. Re-clustering of 18491 monocytes and macrophages at higher resolution revealed 16 clusters (Figure 1A, and Table S1B and S2B). Of note, the cells from Barman et al. contained FACS-sorted bone marrow classical monocytes (Ly6C^hi^), whereas the rest of the studies contained a combination of monocytes and macrophages. Hence, study-wise segregation of UMAP plots allowed us to distinguish bone marrow classical monocyte subsets from the aortic tissue and plaque-associated monocytes and macrophages (Figure 1B). Monocytes and macrophages were identified based on the expression of canonical markers such as *Ly6C2, Ccr2,* and *Adgre1* (encoding F4/80) (Figure 1C). Subsets included neutrophil-like monocytes (NeuMo) (*Mpo, Elane, Prtn3, Ctsg*; cluster 3), which we previously demonstrated arise from GMPs ^24,26^, and MDP-derived moDC-producing monocytes (DCMo) (*Cd209a, Cd74, H2-Aa*; cluster 10) ^24,26^, as well as two subsets of proliferating monocytes (*Mki67, Top2a, Tuba1b, H2ac4, Mxd3*; cluster 4, and *Ube2c, ccnb2, Cdkn3, Cenpa*; cluster 7) ^22,24,25^, and 5 additional monocyte clusters defined as *Sirpb1c^hi^Gngtl°* (*Sirpb1c, Mmp8, vcan*; cluster 2), *Sirpb1c^hi^Gngt^hi^* (*Sirpb1c, Gngt2, Itgal*; cluster 11), interferon inducing cell (IFNIC) (*Ifit3, Rsad2, Isg15*; cluster 8)^2^, *Retnla^hi^Ear2^hi^*(*Retnla, Ear2, Lyz1*; cluster 9), and lipid associated monocytes (*Lgals3, Anxa2, fabp5*; cluster 5) ^2,3,24,25^, (Figure 1A, 1D and S3, and Table S1B and S2B). A *Trem2*^hi^ macrophage (*Trem2*^hi^ Mφ) cluster (*Cd9, Cd72, Spp1, Trem2, Slamf9*; cluster 6) was the only exclusive macrophage subset, whereas chemokine^hi^ monocytes and macrophages (chemokine^hi^ Mo/Mφ) (*Cxcl1, Cxcl2, Ccl4*; cluster 0), tissue-resident Mo/Mφ 1 (*Cd209d, Cd209f, Lyv1, Folr2*; cluster 1), and tissue-resident Mo/Mφ 2 (*Nrh1, Cd51, Fabp4, Fabp5*; cluster 12) shared both monocyte and macrophage gene signatures (Figure 1A, 1D and S3, and Table S1B and S2B) ^2,3^. Cells from three smaller clusters, which were neither monocytes nor macrophages, but showed signature gene expression of vascular endothelial cells (*Cavin2, Cldn5, Ptprb, Podxl*; cluster 13), B cells (*Igha, Igkc, Jchain*; cluster 14), and mixed cells (cluster 15) were excluded from further analysis. Data showing the proportion of each monocyte and macrophage cluster across the studies revealed the abundance of these cell subsets in each sample type (Figure 2A).

**Figure 1.**
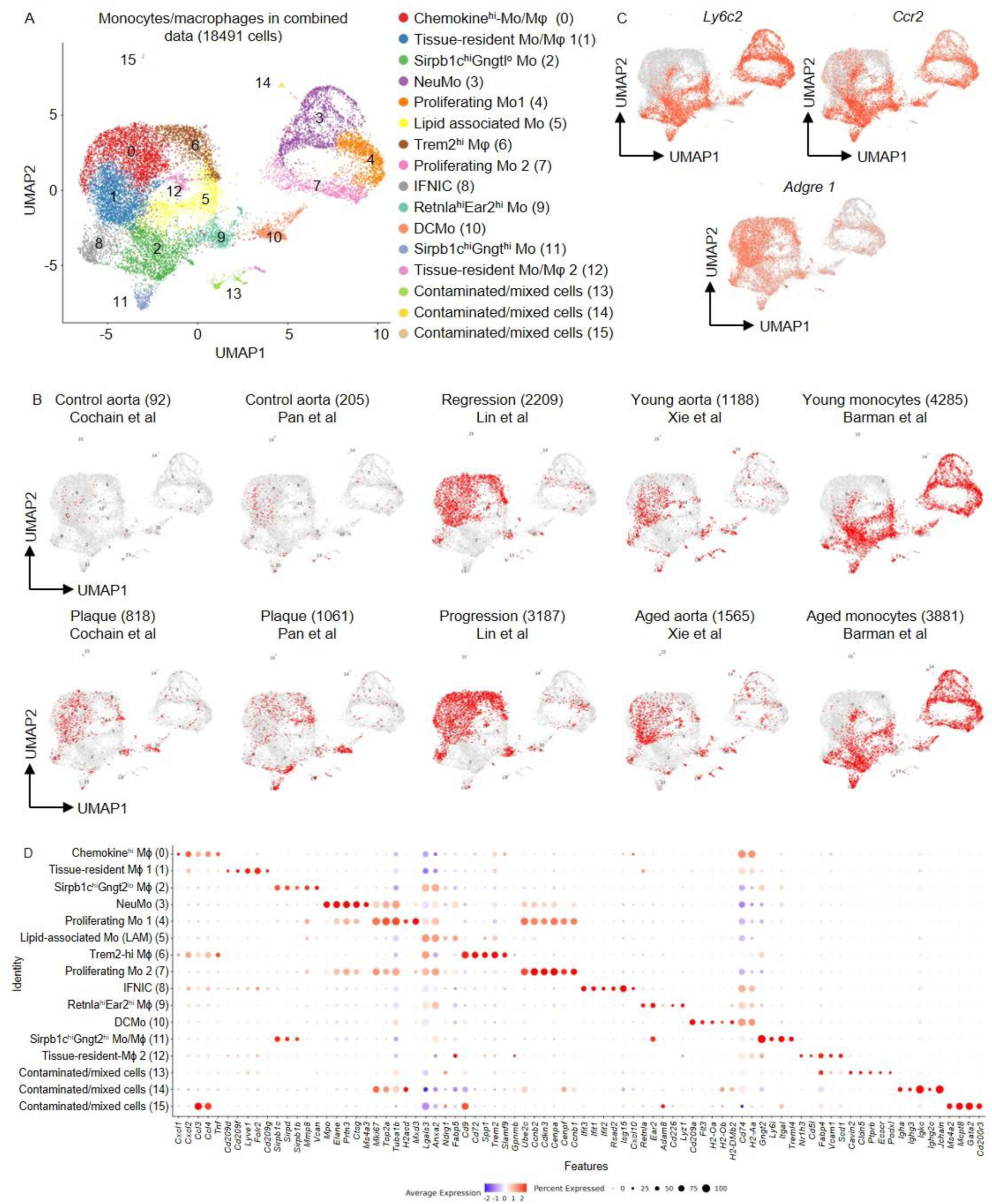
Integrated scRNA-seq analysis reveals monocyte and macrophage subsets in atherosclerotic plaques, aging aorta, and aging bone marrow of mice. (A) UMAP presentation of integrated scRNA-seq gene expression data in 18491 monocytes and macrophages from mouse atherosclerotic plaques, aorta and bone marrow show 16 clusters with major monocyte and macrophage subset identification. (B) UMAP plots according to dataset and experimental conditions show distribution of monocyte and macrophage subsets in each dataset. (C) Expression of monocyte and macrophage maker genes projected onto the UMAP plots. (D) Dot plot shows expression of marker genes across monocyte and macrophage clusters.

**Figure 2.**
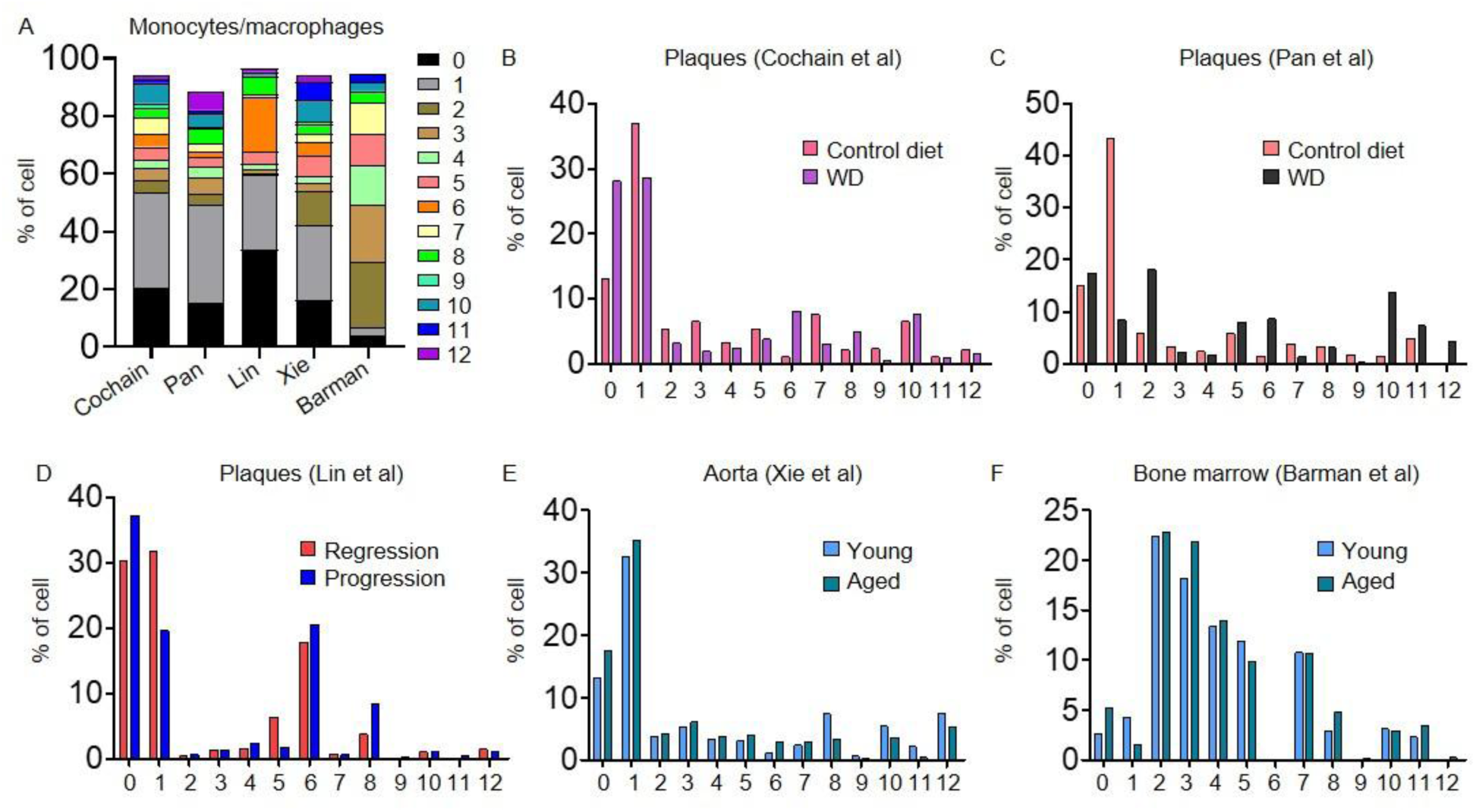
Specific monocyte and macrophage subsets are enriched in atherosclerotic plaques and aging aorta. (A) Bar graph shows proportion of monocyte and macrophage clusters across the studies in the integrated scRNA-seq dataset. (B-F) Proportion of each monocyte and macrophage cluster according to the experimental conditions and models in the integrated scRNA-seq dataset.

Closer examination revealed that the proportion of chemokine^hi^ Mo/Mφ (cluster 0) was higher in atherosclerotic plaques compared to the healthy aorta, and in the aging aorta compared to the young aorta (Figure 2B-2F). These cells were also more abundant in aging bone marrow than in young bone marrow, which indicates that the chemokine^hi^ Mo/Mφ accumulating in the atherosclerotic and aging aortas are of bone marrow origin (Figure 2B-2F). The *Trem2*^hi^ foamy macrophages (cluster 6) were also increased in the plaques and aged aorta, but were negligible in the bone marrow, indicating differentiation of these macrophages locally within the aging and atherosclerotic aorta. On the other hand, the major tissue-resident Mo/Mφ (cluster 1) consistently declined in the plaques, but were unaltered in the aged aorta, indicating that the abundance of these resident macrophages is impacted exclusively by atherosclerosis but not by aging. (Figure 2B-2F). Together, these data suggest that atherosclerosis and aging alter specific aortic macrophage subsets in a similar fashion, but with some differences.

### Atherosclerosis and aging upregulate *Cd74* and MHCII gene expression in monocytes and macrophages

We then interrogated if there are common overlapping genes that are differentially expressed between the monocytes and macrophages from aged bone marrow, aged aorta and atherosclerotic plaques. Analysis revealed 443 DEGs between healthy aorta versus atherosclerotic plaques, 985 DEGs between regressive versus progressive plaques, 443 DEGs between young versus aged aorta, and 153 DEGs between young versus aged bone marrow (Figure 3A, 3B, and Table S3). There were 10 DEGs (*H2-Eb1, H2-Aa, H2-Ab1, Cd74, H2-DMb1, Clec4e, Mt1, Erdr1y, Pros1,* and *Ighm)* that overlapped between all comparisons, and interestingly, five of these genes are associated with MHCII (*H2-Eb1, H2-Aa, H2-Ab1, Cd74,* and *H2-DMb1)* and were found to be consistently upregulated in the monocytes and macrophages from established and progressive atherosclerotic plaques, aged aorta, and aged bone marrow (Figure 3A-3C, and Table S3). These common upregulated genes are markers of MDP-derived DCMos ^24–26^, and were highly expressed in several aortic monocyte and macrophage clusters such as chemokine^hi^-Mφ, Tissue-resident Mφ 1, *Trem2*^hi^ Mφ, IFNIC, *Retnla*^hi^*Ear2*^hi^ Mo/Mφ, and Tissue-resident Mφ 2 (Figure 3D, S4A, and S4B). Therefore, atherosclerosis and aging upregulate *Cd74* and MHCII genes in monocytes and macrophages.

**Figure 3.**
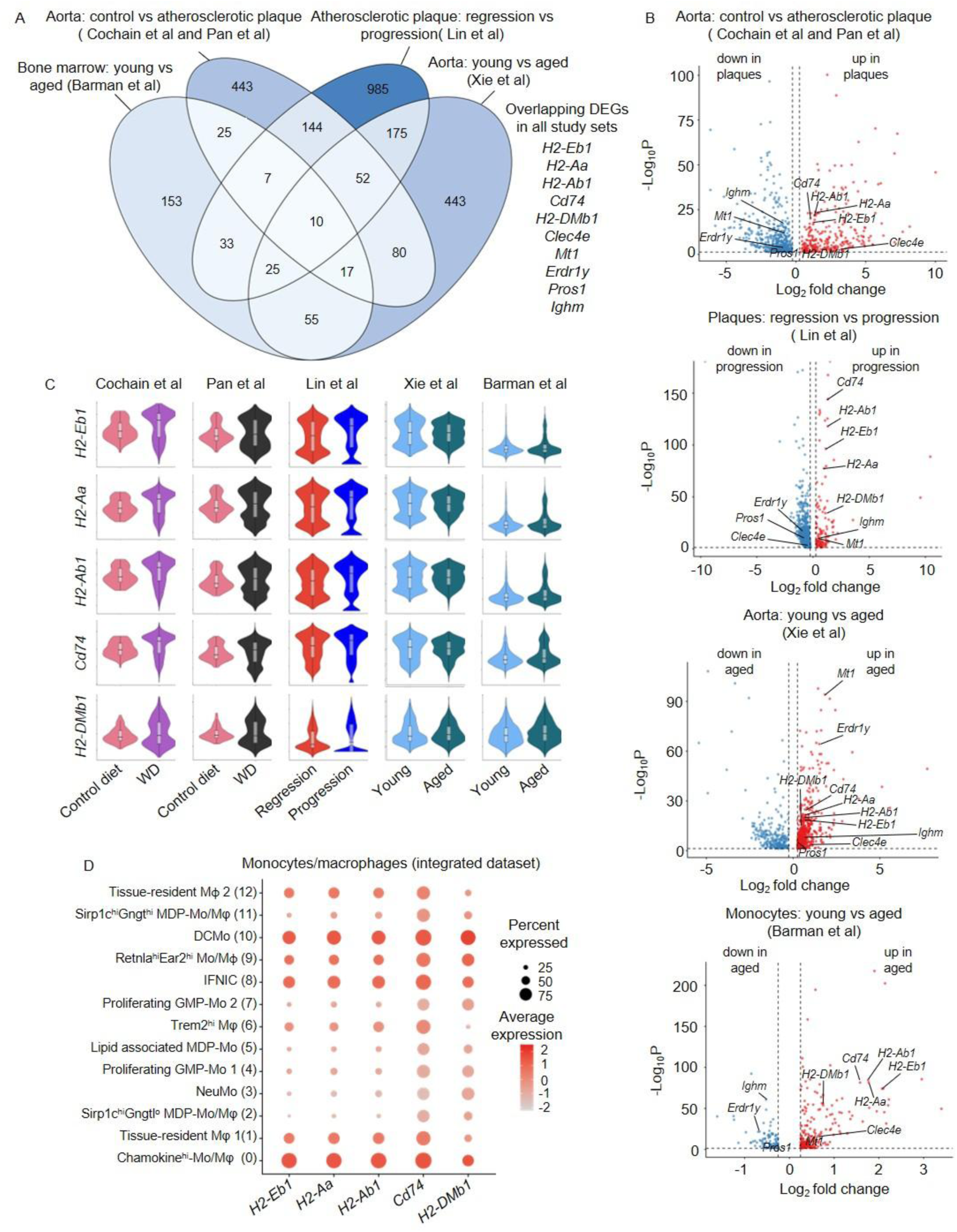
Atherosclerosis and aging upregulate *Cd74* and MHCII gene expression in monocytes and macrophages. (A) Venn diagram shows overlapping DEGs among the data sets. The DEGs were first defined separately by comparing gene expression between the experimental conditions in each dataset. (B) Volcano plot of DEGs that are increased or decreased in atherosclerotic plaques, aged aorta, and aged bone marrow. The common overlapping DEGs are labelled. (C) Violin plots show expression of the common upregulated DEGs between experimental conditions in each dataset. (D) Dot plot shows expression of the common upregulated DEGs across monocyte and macrophage clusters in the integrated dataset.

### Ox-LDL upregulates CD74 expression by mouse macrophages and human monocytes

Previous studies have shown that CD74 expression in atherosclerotic plaques is positively associated with plaque burden, and that CD74 is up-regulated in ox-LDL-induced foamy macrophages, leading to activation of NF-κB and MAPK ^39–41^. The integrated scRNA-seq analysis in the current study also showed upregulated expression of *Cd74* by the monocytes and macrophages from mouse atherosclerotic plaques. Consistent with this, our in vitro experiments showed increased expression of CD74 surface protein in RAW264.7 mouse macrophages and CD74 surface protein and *Cd74* mRNA in THP1 human monocytes by ox-LDL stimulation further validating the induction of CD74 expression in atheroma models of monocytes and macrophages (Figure 4A-4C).

**Figure 4.**
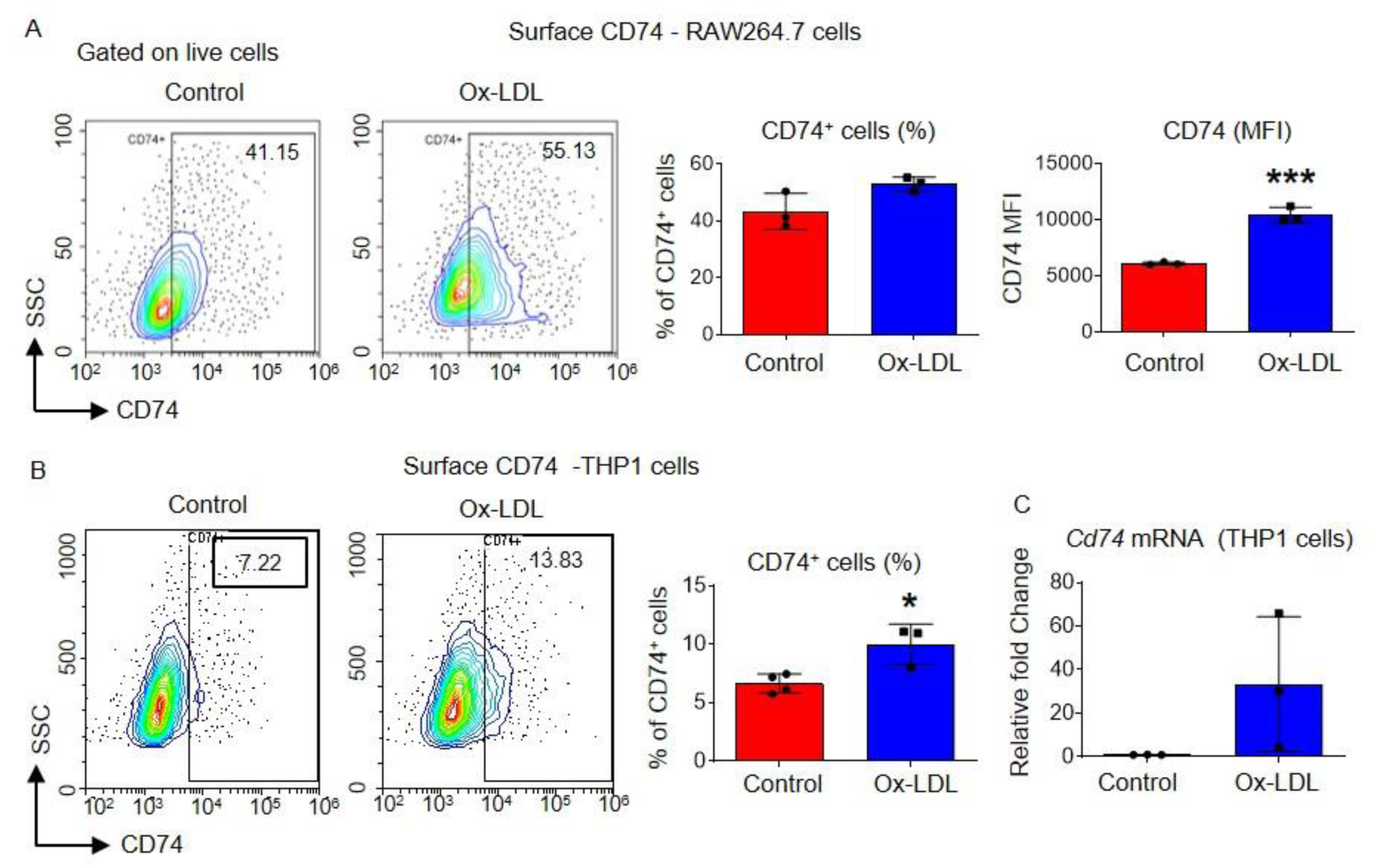
Ox-LDL upregulates CD74 expression by mouse macrophages and human monocytes. (A) CD74 surface protein expression post-ox-LDL (10μg/ml) stimulation was measured in RAW264.7 mouse macrophages by flow cytometry. (B, C) CD74 surface protein (B) and *CD74* mRNA (C) expression in THP1 human monocytes post-ox-LDL (10μg/ml) stimulation was measured by flow cytometry and qRT-PCR respectively. Data are presented as mean plus standard deviation of 3-4 replicates in each group. Statistical significance was assessed by two-tailed Student’s t test (*p < 0.05, ***p < 0.001).

### Aging increases CD74 surface expression on aortic macrophages and bone marrow monocytes in mice

We have previously reported an increased number of CD74^+^ classical monocytes (MDP-derived DCMo) in the bone marrow and blood of aged mice ^24^, and the integrated scRNA-seq analysis in the current study showed increased *Cd74* expression by the monocytes and macrophages in aged mouse aorta. Furthermore, earlier studies have shown that monocytes roll over the endothelial cell layer of tunica intima via weak interactions between monocyte integrins (e.g, VLA-4; very late antigen-4) and vascular endothelial cell adhesion molecules (e.g, VCAM-1; vascular cell adhesion molecule-1) at steady-state. However, increased monocyte integrins can cause stronger monocyte-endothelial cell interaction, leading to monocyte arrest and atherosclerosis progression in response to microinjuries on arterial walls ^1,42^. Hence, we experimentally validated CD74 and VLA-4 expression on aortic macrophages from young and aged mice. Although there was no change in total and VLA-4^+^ macrophages (CD11b^+^F4/80^+^ and CD11b^+^F4/80^+^ VLA-4^+^ cells), their expression of CD74 was significantly increased in the aorta of aged mice in comparison to the young mice (Figure 5A and 5B). We also checked VLA-4 and CD74 expression by bone marrow monocytes from young and aged mice. In contrast to high expression of VLA-4 by the aortic macrophages from both young and aged mice the majority of bone marrow monocytes from young mice were VLA4^-^, and a VLA4^+^ subset emerged in aged mice (Figure 5B and 5C). Importantly, VLA4^+^ monocytes showed higher expression of CD74 when compared with their VLA4^-^ counterparts (Figure 5D). Together, these data indicate that CD74 expression is increased in bone marrow monocytes and aortic macrophages from aging mice, and that increased monocyte and macrophage expression of CD74 is a common phenotype of between atherosclerosis and aging.

**Figure 5.**
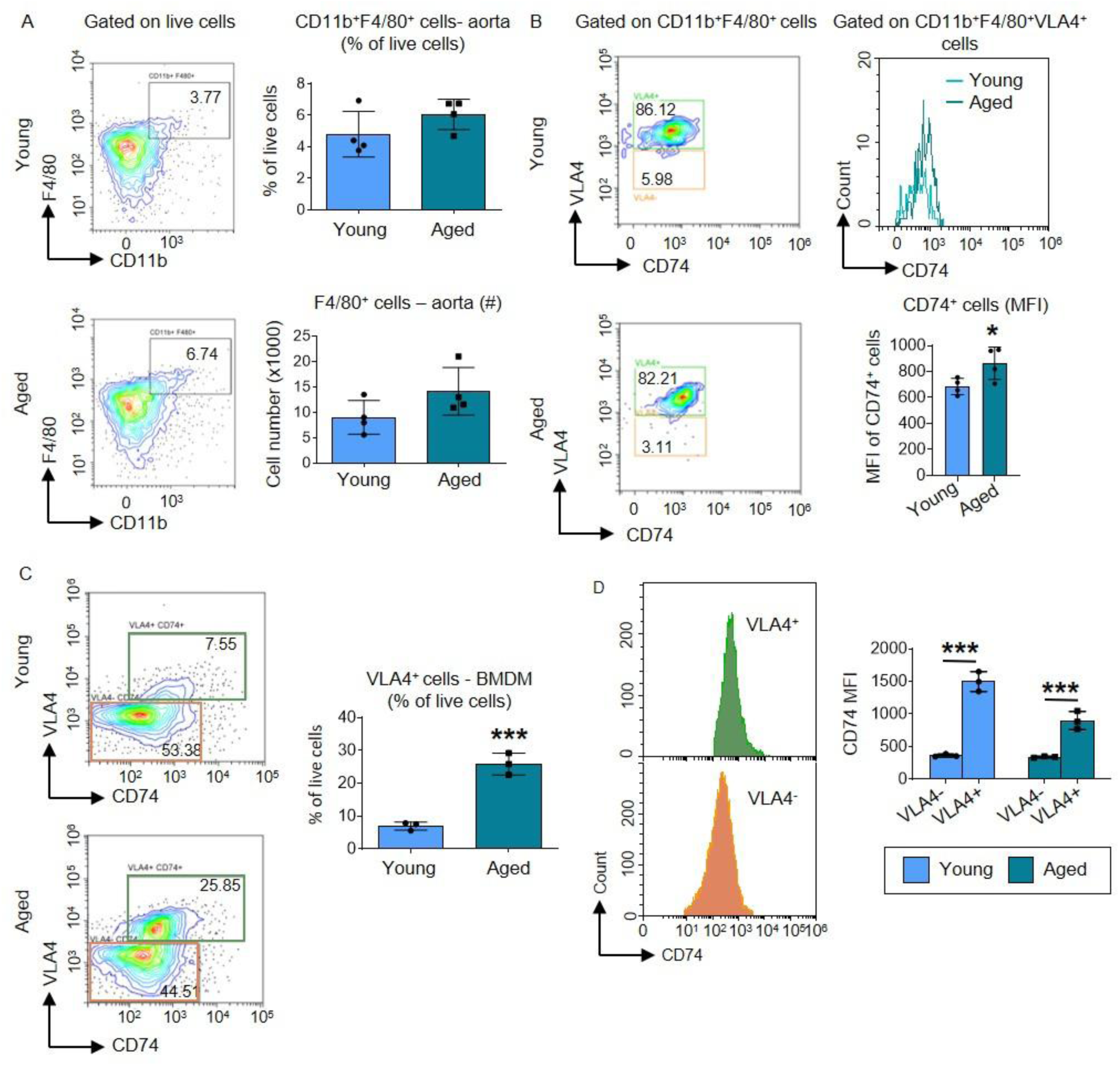
Aging increases CD74 surface expression on aortic macrophages and bone marrow monocytes in mice. (A) Representative flow cytometry plots, percentage and number of CD11b^+^F4/80^+^ macrophages in the aorta of young (3-month old) and aged (26-month old) mice. (B) Representative flow cytometry plots for CD11b^+^F4/80^+^VLA4^+^ macrophages and histogram overlay for CD74 expression and mean fluorescence intensity (MFI) of CD74 on CD11b^+^F4/80^+^ VLA4^+^ macrophages. (C, D) Representative flow cytometry plots (C), and percentage of VLA-4^+^CD74^+^ and VLA-4^+^CD74^-^ (D) in bone marrow monocytes form young and aged mice. Data are presented as mean plus standard deviation of 4 mice in each group (A, B), and 3 replicates of cultures established by pooling cells from 4 mice in each group (C, D). Statistical significance was assessed by two-tailed Student’s t test (*p < 0.05, ***p < 0.001).

### Monocytes from aged mice show increased adhesion to aortic endothelial cells

Monocyte arrest to vascular endothelial cells is a critical step of atherosclerotic plaque development ^1,42^. Therefore, we examined whether aging promotes monocyte adhesion to aortic endothelial cells. Direct reciprocal co-cultures of bone marrow monocytes and AECs from young and aged mice showed that aged CD11b^+^ cells, including monocytes (CD11b^+^CD115^+^ cells), adhered more readily than their young counterparts to both young and aged AECs (Figure 6A, 6B, S5A, and S5B). However, the ability of AECs to arrest monocytes and other CD11b^+^ cells decreased with aging. Upregulated expression of monocyte integrins by the chemokine macrophage migration inhibitory factor (MIF) upon binding to its receptor CD74 is believed to play important roles in monocyte adhesion ^39,40^. Hence, we checked whether ox-LDL plus MIF have a further impact on monocyte adhesion to AECs. We found that although ox-LDL plus MIF marginally decreased monocyte and other CD11b^+^ cell adhesion to young AECs, they incremented these cells’ adhesion to aged AECs (Figure 6A, 6B, S5A, and S5B). Together, these data show that a higher number of aged monocytes adhere to AECs, and that aged AECs have a weaker ability to bind monocytes, which can be reversed by ox-LDL plus MIF stimulation.

**Figure 6.**
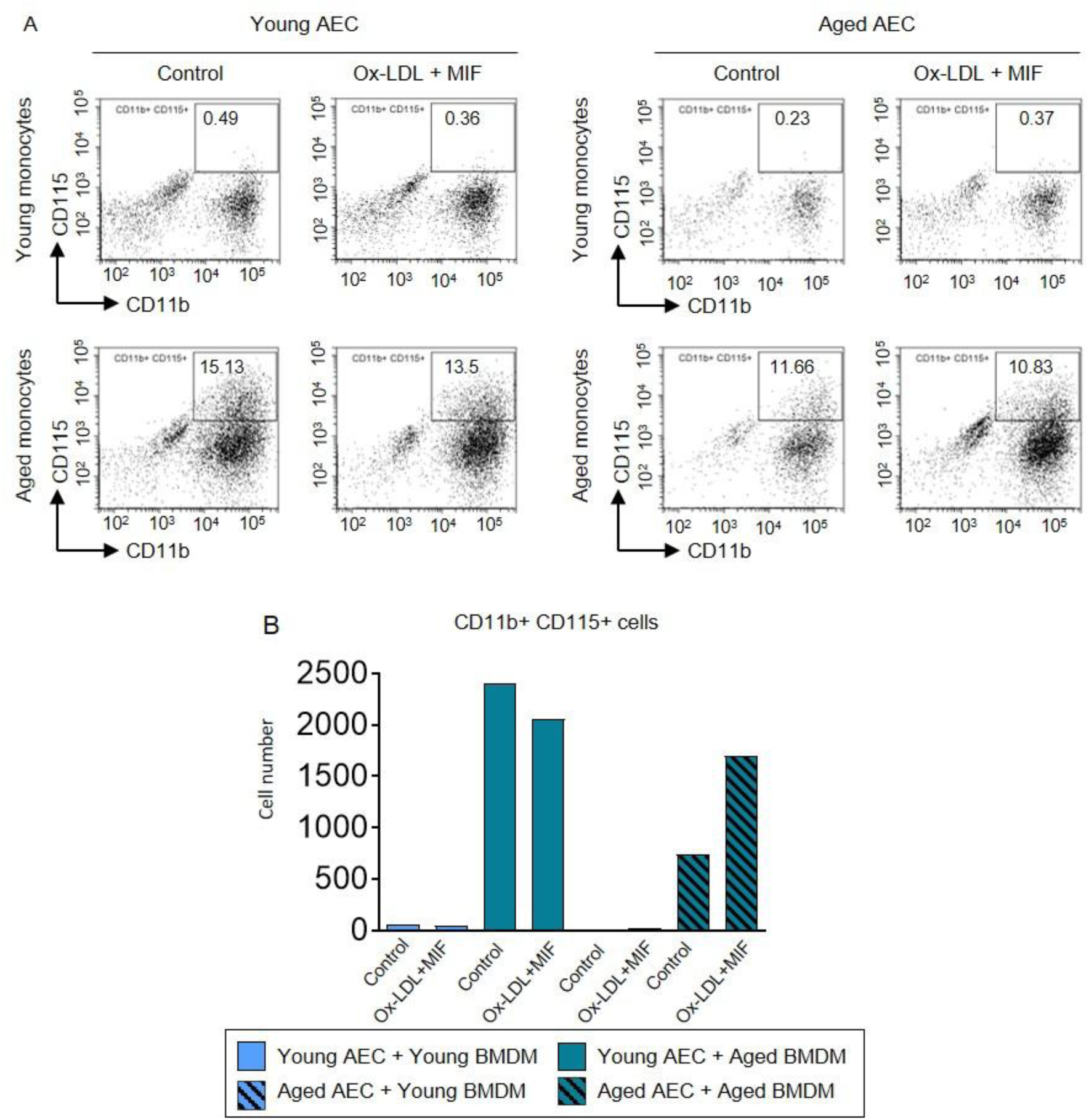
Monocytes from aged mice show increased adhesion to aortic endothelial cells. (A, B) Co-cultures were developed by using aortic endothelial cell monolayer and M-CSF differentiated bone marrow monocytes from young and aged mice. The co-cultures were left untreated or stimulated with ox-LDL (10μg/ml) + MIF (500ng/ml) for 8 hours. Flow cytometry plots (A) and bar graph (B) show difference in the percentage and number of adhered CD11b^+^CD115^+^ cells respectively in the co-cultures. The endothelial cells and monocytes were derived from the cultures of total cells pooled from 4 mice in each group.

## Discussion

Collectively, our data indicate that specific macrophage subsets are elevated in atherosclerotic plaques and aged aorta of mice, implying that these macrophages play a role in connecting atherosclerosis with aging. We also found that MHC class II antigen presentation genes such as *H2-*genes and *Cd74* (which has dual roles as both an MHC II chaperone and as a receptor for MIF) are positively associated with atherosclerosis and aging. In addition, our data show that aortic macrophages in aged mice have higher expression of CD74, and that monocytes from aged mice adhere more to aortic endothelial cells.

Atherosclerotic plaques display heterogeneity in monocytes and macrophages ^2–8^. However, their origin and function are still not completely understood. It is believed that local proliferation of tissue-resident macrophages of both embryonic and postnatal bone marrow origin, and infiltrating monocyte-derived and vascular smooth muscle cell transdifferentiated macrophages contribute to the plaque macrophage pool at different stages of disease progression ^4,6,9,43–46^. For instance, early studies indicated that infiltrating monocyte-derived macrophages are significant participants in developing plaques during early atherosclerosis, whereas proliferation of resident plaque macrophages dominates accumulation of lesional macrophages during the late stage of atherosclerosis in mice ^45^. Later studies showed that resident macrophages, including aortic intimal-resident macrophages (Mac^AIR^) transform into foamy cells contributing to early lesions, and promote infiltration of circulating monocytes to the plaques ^6,47^. Hence, bone marrow monocytes, which are elevated by atherosclerosis-induced monopoiesis, play critical roles in atheroma progression^43,44^. Ontogeny also dictates recruitment of monocytes to different tissues: while both GMP-and MDP-derived monocytes infiltrate the intestine and lung, GMP-derived monocytes exclusively give rise to meningeal dura mater macrophages ^25^. Furthermore, monocyte production via GMPs or MDPs is differentially influenced by distinct microbial stimuli, and we have previously shown that while aging increases monopoiesis by both GMPs and MDPs in mice, there is a notably strong increase in MDP-derived DCMo production ^24–26^. However, it’s unclear whether atherosclerosis-induced monopoiesis involves one or both of these progenitors. Our finding of elevated macrophages with MDP lineage features (increased expression of *Cd74* and *H2-*genes) in atherosclerotic and aged aorta may imply that atherosclerosis and aging promote recruitment of MDP lineage monocytes to the aortic intima, where they differentiate into plaque macrophages. Future studies using lineage-tracing mouse models of aging and atherosclerosis will be helpful to define the ontogeny of aortic macrophages.

Atherosclerotic plaque macrophages also show functional diversity ^3,5,9^. For instance, chemokine^hi^ macrophages, which constitute the highest proportion within the plaque macrophages, are pro-atherogenic in nature due to their ability to drive inflammation similar to M1 macrophages ^3,5^. In contrast, TREM2^hi^ foamy macrophages play important roles in atheroprotection by reducing the necrotic core area via increased lipid uptake, efferocytosis, and inhibition of foamy macrophage death ^10,11^. Lyve1^+^ resident-like macrophages maintain tissue homeostasis and inhibit plaque progression by controlling collagen production by SMCs, promoting endocytosis, and a repair mechanism similar to M2 macrophages ^3,9,12,13^. Our findings of increased proportion of chemokine^hi^ inflammatory Mo/Mφ and *Trem2*^hi^ foamy macrophages in atherosclerotic plaques and aged aorta further support the notion that altered macrophage functions in the aged aorta play important roles in atherosclerosis progression.

Aging causes a slow decline in vascular health and the immune system, leading to endothelial cell and monocyte dysfunction ^48^. Yet, the factors that underlie aging-associated risk of atherosclerosis are unclear ^49^. Our data showing adhesion of a higher number of monocytes to aortic endothelial cells, along with upregulated expression of CD74 on aortic macrophages and VLA4^+^ bone marrow monocytes in aged mice, may imply that CD74-mediated adhesion and transmigration of monocytes to the subendothelial layer of the aorta is critical for inducing a pre-atherosclerotic condition in aging ^39,50^. Furthermore, it is interesting that MIF restored the ability of aged AECs to adhere monocytes. It will be interesting to see if a similar mechanism underlies this in vivo, leading to monocyte influx into the aging mouse aorta.

Increased expression of *Cd74* and *H2* genes in monocytes and macrophages from atherosclerotic plaques, aged aorta, aged bone marrow may indicate that antigen presentation by MHC II is impacted by both atherosclerosis and aging. In this context, a recent study has shown that increased antigen presentation by MHC II is associated with atherosclerosis progression, and that MHC II overexpression leads to exacerbated atherosclerosis ^51^. In contrast to this, MHCII deficiency in ApoE^−/−^ mice has shown increased atherosclerosis due to reduced number of regulatory T cells, suggesting a protective role of MHCII-mediated antigen presentation in atherosclerosis ^52^. Similarly, recently, aging-associated DNA damage and dysregulated autophagy in macrophages have been shown to induce autoantigen presentation on MHC II in mice ^53^. It will be important to study how antigen presentation by MHC II connects atherosclerosis with aging.

In conclusion, the current study reveals that *Cd74* and *H2* genes (MDP lineage signature) are upregulated in aortic macrophages from atherosclerotic and aged mice. Recently, VCAM-1 and VLA-4 inhibitors have emerged as promising drug targets for treating atherosclerosis by reducing macrophage recruitment to the arteries ^54,55^. However, these strategies are limited by the inhibition of pan-monocyte rolling, causing toxicity. Future studies to investigate atheroprone monocytes in aging will help designing therapeutic strategies to prevent atherosclerosis by blocking recruitment of these monocytes to the arteries.

## Materials and Methods

### Mice

Wild-type C57BL/6 mice were purchased from The Jackson Laboratories and maintained at the Centre for DNA Fingerprinting and Diagnostics (CDFD) animal facility. Young (3-month-old) and old (26-month-old) male mice were used for the experiments. Committee for Control and Supervision of Experiments on Animals (CCSEA), India regulations were followed to perform all procedures. Mice were fed a regular chow diet *ad libitum*.

### Cell Culture

RAW264.7 and THP1 cells were cultured in DMEM and RPMI, respectively, supplemented with FBS (10% v/v), penicillin (50 U/ml), streptomycin (50 U/ml), and L-glutamine (2 mM). Cells were stimulated with ox-LDL (10 ug/mL) for 6-24 hours and used for qRT-PCR or flow cytometry analysis.

### Mouse bone marrow monocyte culture

Total bone marrow cells were collected by high-speed centrifugation of the tibias and femurs from both hind legs ^56^. RBCs were lysed, and the cells were cultured in the presence of recombinant mouse M-CSF (50 ng/ml) for 3 days to induce monocyte differentiation. After 3 days, the non-adherent monocytes were collected for the experiments.

### Mouse AEC culture

The whole thoracic aortae were cut into small pieces and digested with collagenase type 2. The single cells from collagenase-digested aortae were cultured in human large vessel endothelial cell basal medium supplemented with low serum growth for three days. The adherent endothelial cells were pooled for the experiments.

### Co-culture Assay

1x10^5^ AECs were cultured per well in a 96-well plate overnight for making a monolayer. 1x10^5^ bone marrow monocytes were seeded on top of the AEC monolayer, followed by stimulation with ox-LDL (10 ug/mL) plus MIF (500 ng/mL) or untreated for 8 hours. After PBS washing for three times, the adherent cells were detached for flow cytometry analysis.

### Flow Cytometry

Antibodies used for flow cytometry are listed in Table S4. To assess aortic macrophages, cells were stained with antibodies against Zombie-R718, CD11b-PE/Dazzle 594, F4/80-PE, VLA4-FITC, and CD74-AF647. For bone marrow monocytes, cells were stained with antibodies against VLA4-FITC and CD74-AF647; THP1 and RAW264.7 cells were stained with the antibody against CD74-AF647. Cells were incubated with blocking buffer (1X PBS, 0.5% BSA, 1% FBS) before staining to prevent non-specific antibody binding. Flow cytometry was performed using a CytoFlex (Beckman Coulter) and data were analyzed with CytExpert 2.6.

### qRT-PCR

RNAisoplus (Takara) was used to isolate total RNA, and PrimeScript™ 1st strand cDNA Synthesis Kit (Takara) was used to prepare first-strand cDNA using the manufacturer’s protocol. Custom-designed primers and TB green master mix (Takara) were used to perform qRT-PCR. Relative gene expression of *Cd74* (mouse; forward 5’gctccacctaaagtactgacc3’ and reverse 5’agttaccgttctcgtcgca3’, and human; forward 5’ctccaccgaaagtactgacc3’ and reverse 5’agatagttgccgttctcgtcg3’) was measured using *Rps29* (forward 5’ ggtcaccagcagctctactg3’ and reverse 5’gtccaacttaatgaagcctatgtcc3’) and beta-2-microglobulin (B2M) (forward 5’ agatgagtatgcctgccgtg3’ and reverse 5’gcggcatcttcaaacctcca3’) as reference transcripts for mouse and human, respectively.

### Integrated scRNA-seq analysis

Publicly available mouse scRNA-seq datasets were retrieved from the Sequence Read Archive (SRA) ^3–5,24,38^. For datasets lacking standard barcode and UMI fastq files, processed BAM files were downloaded and converted to FASTQ formats using the 10x Genomics bamtofastq utility. Raw sequencing reads were aligned to the mouse reference genome and quantified using STARsolo. Cell filtering and background noise removal were performed natively during alignment using the EmptyDrops CR algorithm. Appropriate 10x Genomics whitelists (v2/v3) and UMI lengths were dynamically applied based on the specific chemistry utilized by the original authors.

Downstream computational analysis was performed in R (version 4.4.3) using Seurat (version 5.4.0). Raw count matrices were loaded, and Ensembl IDs were mapped to gene symbols. To exclude low-quality cells, empty droplets, and multiplets, adaptive quality control thresholds were applied to account for the differential capture sensitivities of the 10x Genomics Chromium Single Cell 3’ v2 and v3 chemistries utilized across the studies. While a universal minimum threshold of 200 features per cell was applied to all samples, the upper feature limits were dynamically adjusted because v3 chemistry inherently yields higher gene detection rates, and applying standard v2 cutoffs would improperly discard healthy cells. The maximum feature counts were capped at 9000 genes for the v3-profiled Xie et al. dataset, whereas a stringent 5000 gene upper limit was set for the v2-profiled datasets such as Cochain et al., Lin et al., and Pan et al. Barman et al. datasets, which were processed with a mixture of v2 and v3 chemistries, were assigned appropriate upper limits. The maximum allowable mitochondrial read fractions were restricted to < 10% for all aortic datasets, but permitted up to < 15% for the Barman et al. bone marrow samples to accommodate tissue-specific metabolic baselines. Following filtration, log-normalization was performed in the retained cells, and the top 2000 highly variable features for each dataset were identified using the variance-stabilizing transformation (vst) method.

To remove technical batch effects arising from varying chemistries and laboratory origins, Canonical Correlation Analysis (CCA) was applied. Integration anchors were identified using FindIntegrationAnchors based on the consensus variable features (SelectIntegrationFeatures), followed by dataset merging via IntegrateData. The integrated matrix was scaled, and Principal Component Analysis (PCA) was performed using the first 30 principal components. Uniform Manifold Approximation and Projection (UMAP) and Louvain clustering (resolution 0.5) were executed using a fixed random seed (seed = 42) to guarantee topological reproducibility. Clusters were annotated based on canonical marker gene expression.

Differentially expressed genes (DEGs) were evaluated by layer unification (JoinLayers). To remove confounding batch effects in pooled comparisons (e.g., pooled atherosclerosis vs. control, or pooled aged vs. young), logistic regression (test.use = “LR”) was used to regress out “Study” or “Chemistry” as latent variables. Standard Wilcoxon rank-sum tests were used for direct, intra-study comparisons. Significant DEGs were defined by a Benjamini-Hochberg-adjusted p-value < 0.05 and an absolute log2 fold-change ≥ 0.25. Data manipulation was handled using dplyr (v1.2.0), tidyr (v1.3.2), purrr (v1.2.1), and MatrixGenerics (v1.18.1). Data visualizations such as feature plots, violin plots, and dot plots were generated using ggplot2 (v4.0.2), patchwork (v1.3.2), and ggVennDiagram (v1.5.7).

## Statistical analysis

Statistical analysis of flow cytometry data and mRNA expression data was performed using Student’s t-test in Prism 10.3.0 software (GraphPad Inc.), and differences with p ≤ 0.05 were considered significant. Statistical analyses of integrated scRNA-seq data were described above.

## Data availability

The scripts for scRNA-seq analyses will be provided upon request to the corresponding author.

## Acknowledgements

The study was supported by Science and Engineering Research Board/Anusandhan National Research Foundation (SERB/ANRF), Govt. of India grant SRG/2023/001276 and funds from Mahindra University to PKB, and Department of Biotechnology (DBT), Govt. of India BioCARe grant BT/PR50764/BIC/101/1315/2023 to SB. The authors are also grateful for the assistance of the high-performance computing (HPC) cluster unit of Mahindra University and the animal house facility of CDFD.

## Authorship Contributions

AM and PKB designed the project; AM performed the mouse and in vitro experiments and analyses and SK, SB and KG assisted her. ISD performed the scRNA-seq analysis with assistance of SS, and BR and PKB guided them. HSG and TH assisted with the data analysis, interpretation and presentation. AM and PKB wrote the manuscript; all authors edited and/or approved the manuscript.

## Abbreviations

ox-LDL: oxidized LDL
GMP: granulocyte-monocyte progenitor
MDP: monocyte-dendritic cell progenitor
MoDC: monocyte-derived dendritic cells
scRNA-seq: single-cell RNA sequencing
DEG: differentially expressed gene
UMAP: uniform manifold approximation and projection
DC: dendritic cell
NeuMo: neutrophil-like monocytes
DCMo: moDC-producing monocytes
IFNIC: interferon inducing cell
Mo/Mφ: monocytes and macrophages
VLA-4: very late antigen-4
VCAM-1: vascular cell adhesion molecule-1
AEC: aortic endothelial cell
MIF: macrophage migration inhibitory factor
Mac^AIR^: aortic intimal-resident macrophages

