## Supplemental figures for "Increased expression of *Cd74* and MHC II genes by aortic macrophages links atherosclerosis with aging in mice"

### Supplementary figures

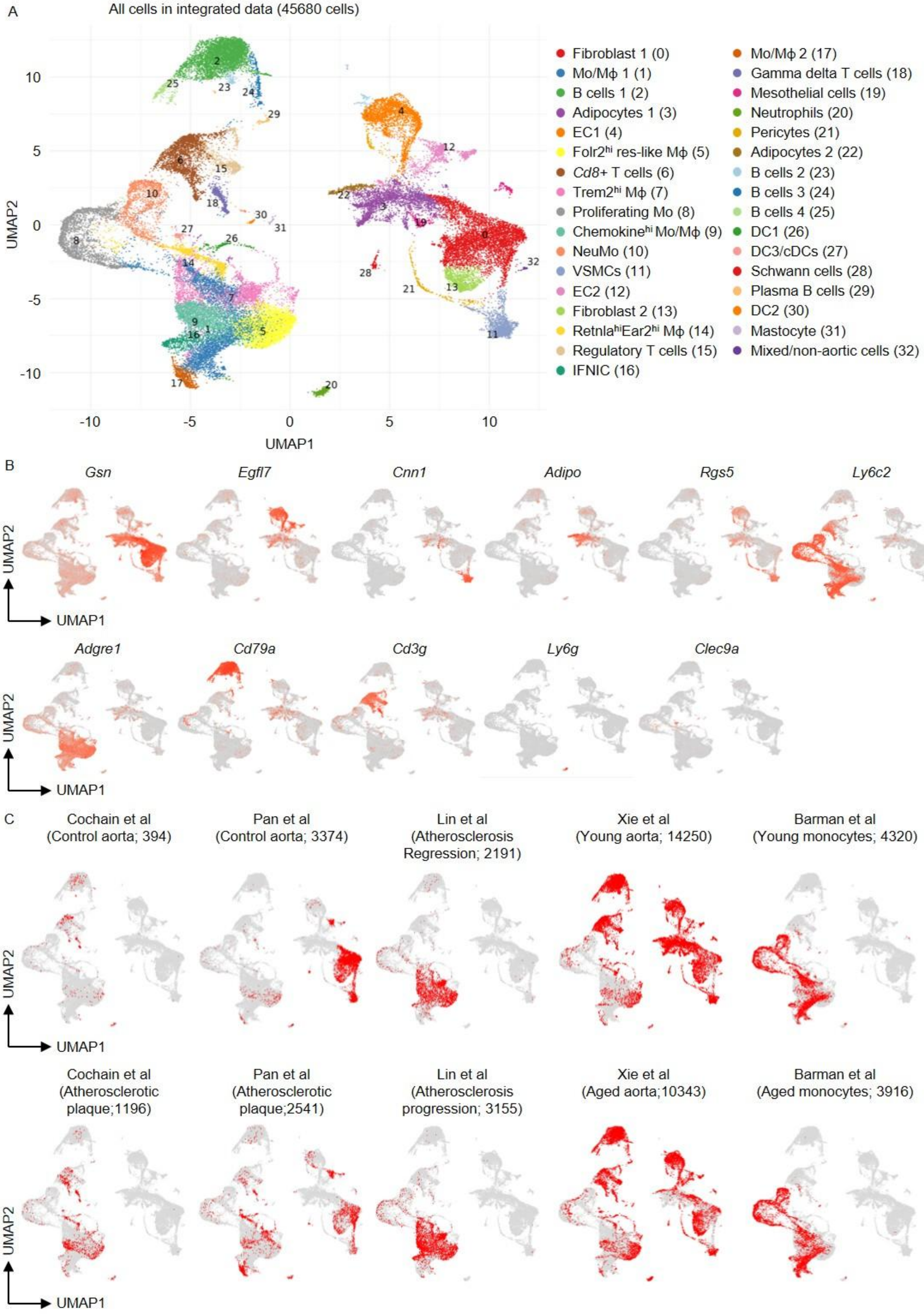

**Figure S1. Integrated scRNASeq analysis of mouse bone marrow, aorta, and atherosclerotic plaques from different experimental conditions**. (A-C) scRNASeq data from Cochain et al. 2018, Pan et al. 2020, Lin et al. 2019, Xie et al. 2022, and Barman et al. 2022 were integrated for the analyses. UMAP representation show 33 clusters. Fibroblasts (*Gsn, Dcn, Serpinf1)*, endothelial cells (ECs) (*Pecam1, Egfl7)* and adipocytes (*Adipoq, Cidec)* each comprised two subsets with enriched expression of *F3 and Fbln7* (fibroblast 1; cluster 0), *Dact2, Sema3e* (fibroblast 2; cluster 13), *Fabp4, Car4* and *Rbp7* (EC1; cluster 4), *Vwf, Plvap* and *Lrg1* (EC2; cluster 12), *Cyp2e1* (adipocytes 1; cluster 3), and *Ces1f and Adrb3* (adipocytes 2; cluster 22). Vascular smooth muscle cells (VSMCs) (*Cnn1, Tagln, Acta2, Tpm2*; cluster 11), mesothelial cells (*Muc16, Msln, Krt19, Dmkn*; cluster 19), pericytes (*Higd1b, Or51e1, Epas1*; cluster 21), Schwan cells (*Mpz, Kcna1, Plp1*; cluster 28) and mastocytes (*Cma1, Mcpt4, Tpsb2*; cluster 31) did not show any subcluster. Within lymphoid lineage, B cells were detected by the expression of *Cd79a* and *Cd79b*, and contained four mature B cell subsets distinguished by differential expression of *Ighd, Fcmr, Pax5, Cd19, Cd22 H2-Ob, Cxcr5, Il4i1, Vpreb3*, and *Tnfrsf13c* (clusters 2, 23, 24 and 25), and one plasma B cell subset (Jchain, Igha, Igkc; cluster 29). T cells (*Cd3g, Trbc2*) consisted of *Cd8*$^{+}$ T cells (*Cd8a, Cd8b1, Nkg7*; cluster 6), regulatory T cells (*Foxp3, Ctla4, Tnfrsf4, Icos*; cluster 15), and gamma delta T cells (*Trgc1, Trdc, Cxcr6*; cluster 18). Myeloid compartment consisted of three DC (detected by *Flt3*) subsets expressing *Tmem150, H2-M2* and *Fscn1* (cluster 26), *Cd209a* and *siglech* (cluster 30), and *Xcr1, Clec9a* and *Naaa* (cluster 27), one neutrophil subset (*S100a8, S100a9, Retnlg, Ly6g*; cluster 20), and nine monocyte and macrophage subsets. Cluster annotation was done by using canonical marker genes from the respective studies and top differentially expressed genes (DEGs) from the current analysis (A). Expression of representative genes of the major cell types projected onto the UMAP plots (B). Projection of single cells in the UMAP space according to dataset and experimental conditions (C).

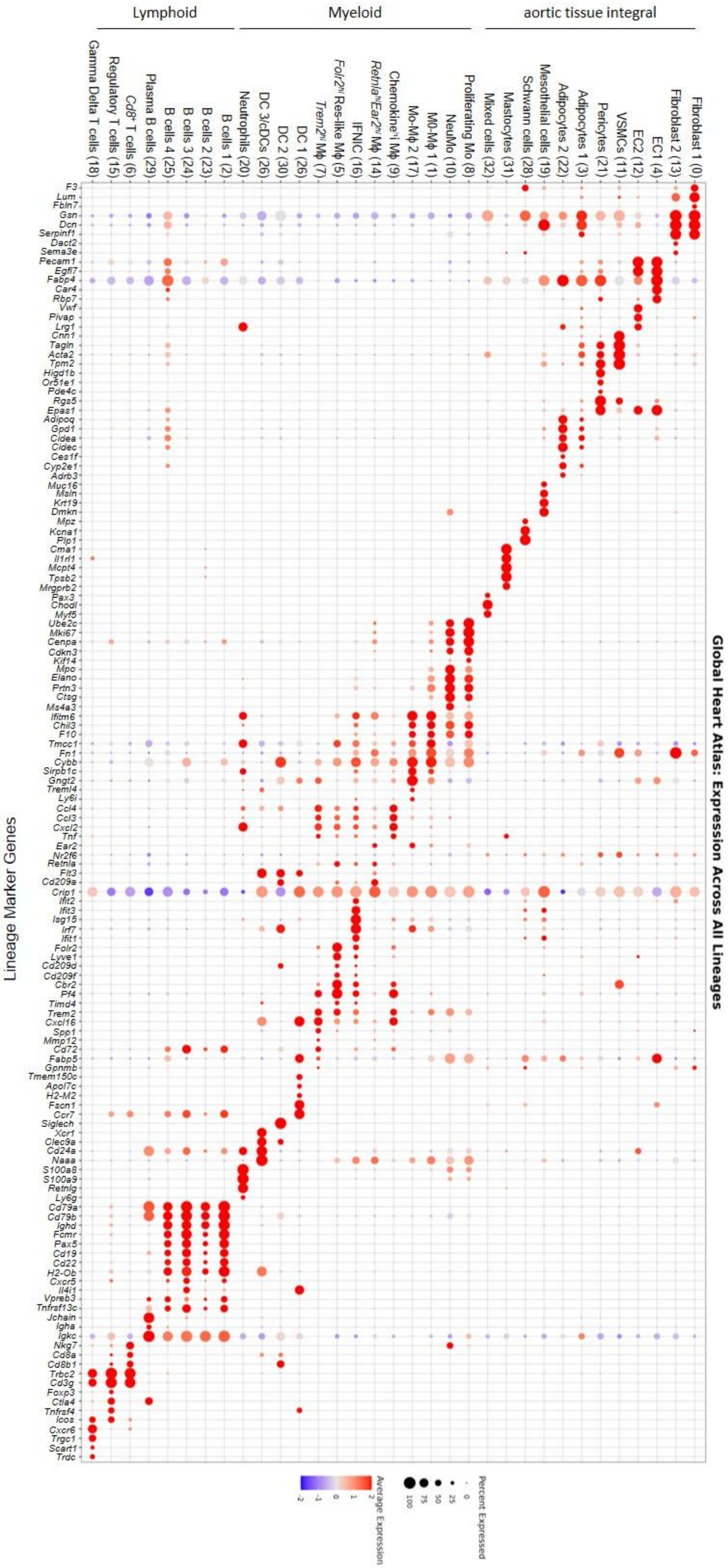


**Figure S2.** Dot plot shows expression of marker genes across the clusters in the integrated scRNAseq data.

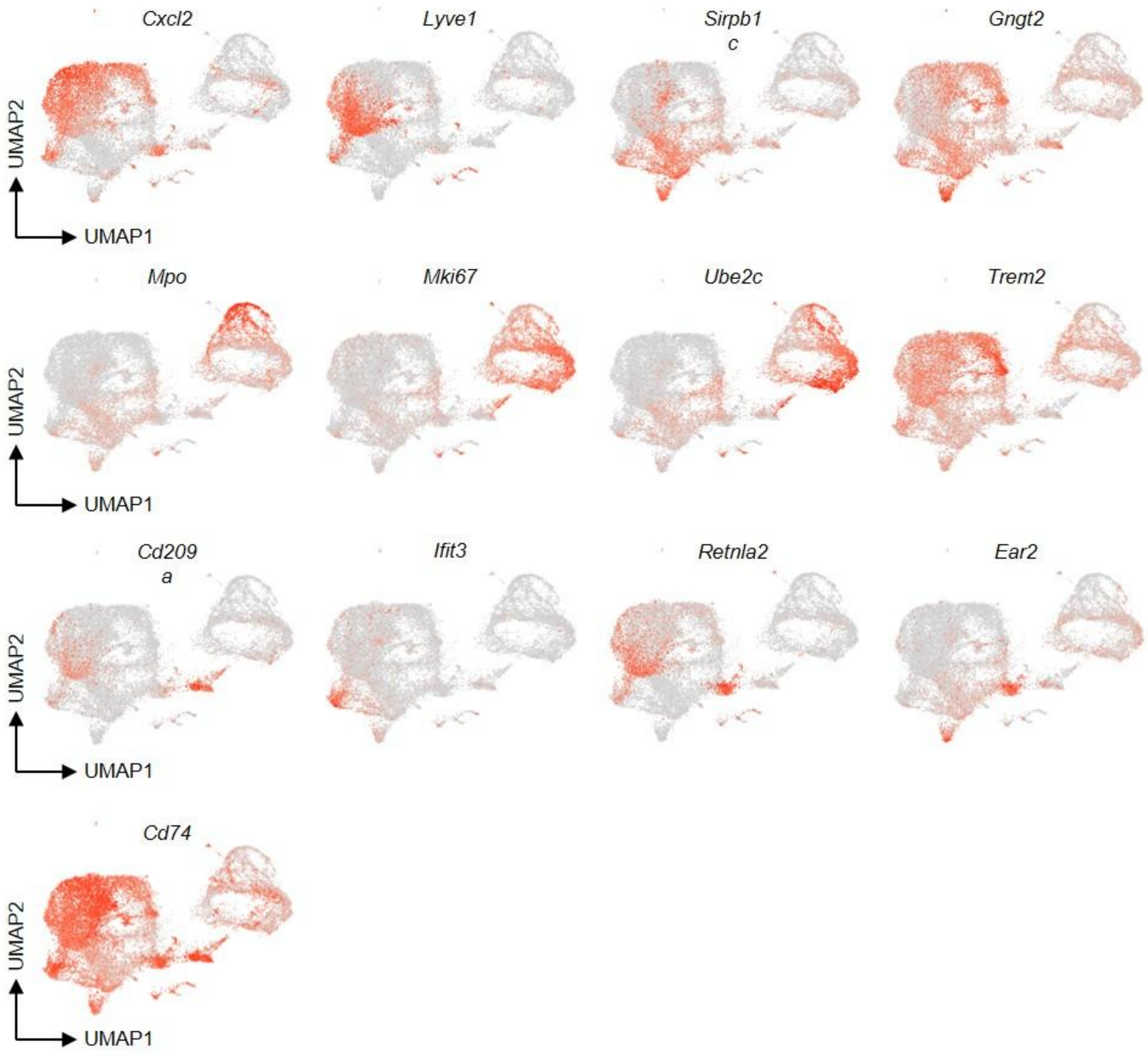


**Figure S3.** Expression of representative genes of the monocyte and macrophage subsets projected onto the UMAP plots.

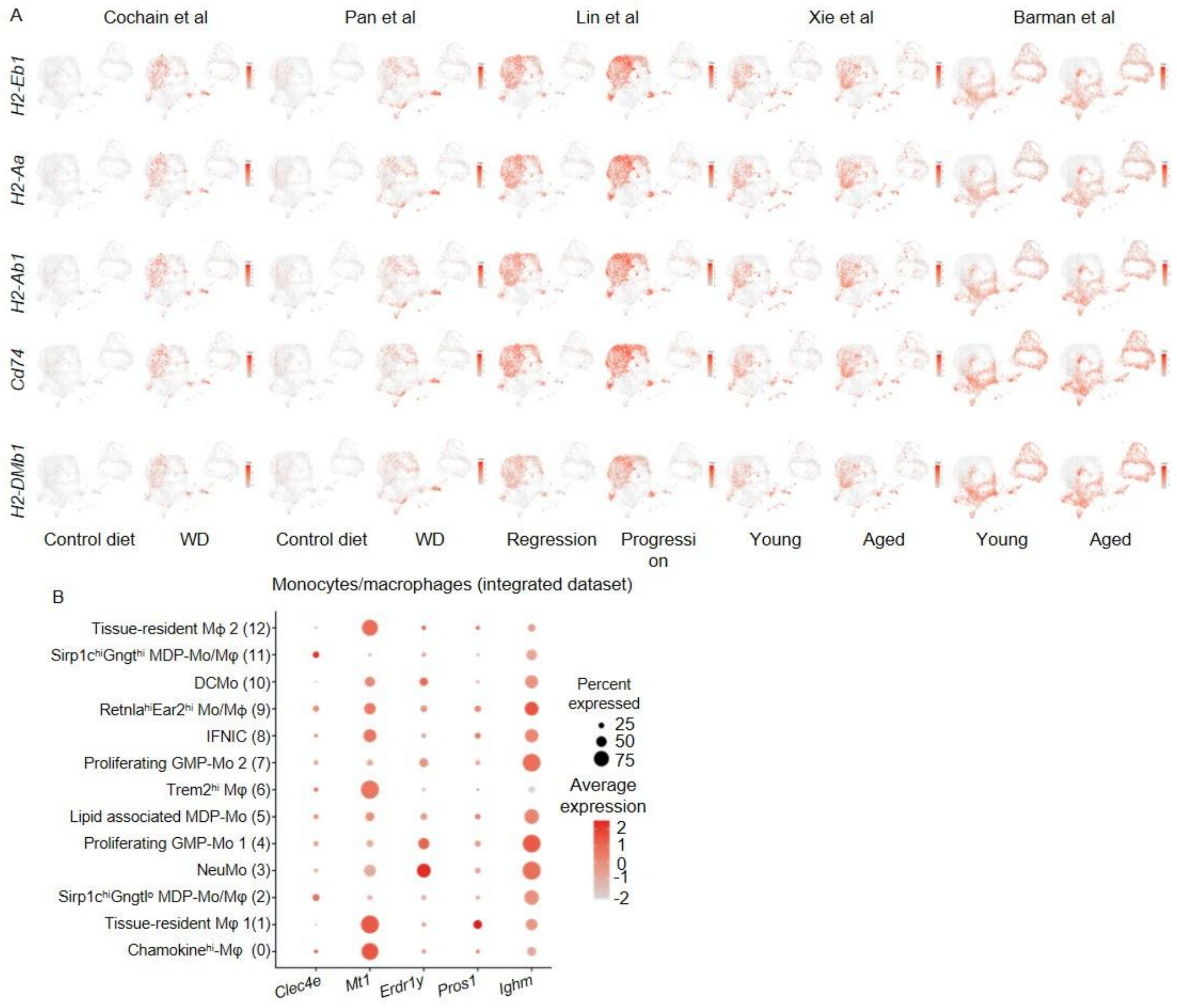


**Figure S4.** (A) UMAP plots show expression of the common upregulated DEGs between experimental conditions in each dataset. (B) Dot plot shows expression of the common DEGs across monocyte and macrophage clusters in the integrated dataset.

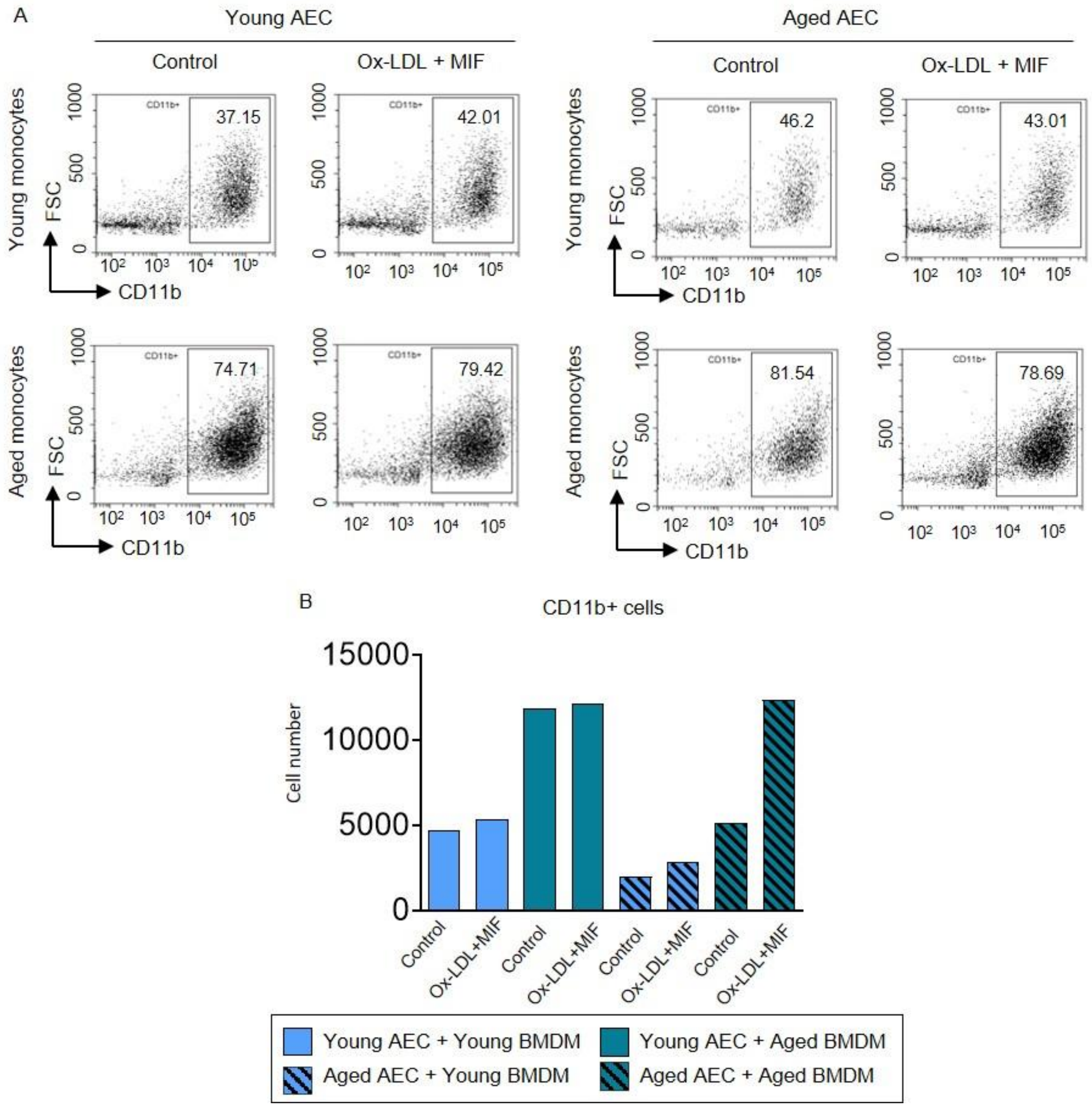


**Figure S5.** (A, B) Co-cultures were developed by using aortic endothelial cell monolayer and M-CSF differentiated bone marrow monocytes from young and aged mice. The co-cultures were left untreated or stimulated with ox-LDL (10µg/ml) + MIF (500ng/ml) for 8 hours. Flow cytometry plots (A) and bar graph (B) show difference in the percentage and number of adhered $CD11b^+$ cells respectively in the co-cultures. The endothelial cells and monocytes were derived from the cultures of total cells pooled from 4 mice in each group.
